# Bacterial *PxRdl2* dsRNA increased the insecticidal activities of GABAR-targeting compounds against *Plutella xylostella*

**DOI:** 10.1101/2021.03.24.436736

**Authors:** Ben-Jie Li, Kun-Kun Wang, Ye Yu, Jia-Qi Wei, Jian Zhu, Jia-Li Wang, Fei Lin, Han-Hong Xu

**Author notes:** **Corresponding authors:** E-mail address (H.H. Xu); (F. Lin). These authors contributed equally to the work (Ben-Jie Li, Kun-Kun Wang).

## Abstract

The utilization of RNA interference (RNAi) for pest management has garnered global interest. The bioassay results suggested the knockout of *PxRdl2* significantly increased the insecticidal activities of the *γ*-aminobutyric acid receptor (GABAR) targeting compounds (Fipronil, two pyrazoloquinazolines and two isoxazolines), thereby presenting a viable target gene for RNAi-mediated pest control. Consequently, we suggest enhancing the insecticidal activities of GABAR-targeting compounds by knockdown the transcript level of *PxRdl2*. Furthermore, *PxRdl2* dsRNA was expressed in HT115 *Escherichia coli* to reduce costs and protect dsRNA against degradation. In comparison to *in vitro* synthesized dsRNA, the recombinant bacteria (ds-B) exhibited superior interference efficiencies and greater stability when exposed to UV irradiation. Collectively, our results provide a new strategy of insecticide spray which combined synergistically with insecticidal activities by suppressing *PxRdl2* using ds-B, and may be beneficial for reducing the usage of insecticide and slowing pest resistance.

## 1. Introduction

Since RNA interference (RNAi) was first discovered in the nematode *Caenorhabditis elegans* (Fire *et al*., 1998), it had been widely used as a powerful genetic tool for research on various taxa, including plants, insects, nematodes, bacteria, fungi, etc. (Koch and Kogel, 2014; Zotti et al., 2018; Vogel et al., 2019). RNAi had been considered as a new generation of pest control technology with great advantages such as safety and fast environmental degradation (Hernández-Soto and Chacón-Cerdas., 2021). However, an efficient double-stranded RNA (dsRNA) delivery system was critical for validating the efficacy of gene silencing (Lu et al., 2023). Although the microinjection was highly efficient, it was difficult to apply in fields because of the high costs and complexity of preparing dsRNA. Fortunately, ingestion of specific dsRNA through the oral cavity can trigger a silencing response in most insect tissues, which paves the way for the development of RNAi-based agricultural pest control if dsRNA can be protected from degradation in the environment and in the insect gut by suitable delivery methods (Caccia et al., 2020).

Currently, combining dsRNA with nanomaterials or expressing dsRNA in bacteria are the two main methods of dsRNA-insect delivery. It has been reported that nanomaterials can improve the efficiency of delivery allowing for dsRNA to permeate cell membranes directly or by endocytosis (Al-Jamal et al., 2011); thus, this material has become a new type of the delivery agent and has been used in pest RNAi control (Sun et al., 2020). Employing bacteria as delivery vectors of dsRNA molecules is another approach which was first carried out in *C. elegans* (Timmons et al., 2001). Considering the technology and economic-related issues of dsRNA synthesis *in vitro*, many researchers preferred to use dsRNA that expressed in bacteria to induce insect RNAi to control pests (Bento et al., 2020; Chikami et al., 2021; Dhandapani et al., 2020). Except the efficiency of dsRNA delivery, other limitations related to the application of dsRNA for pest control include the identification of appropriate target genes and increasing insect lethality. From these points, genes targeted by chemical insecticides and bio-insecticides might be ideal sites for dsRNA targeting.

With the cultivation of cruciferous crops, the *Plutella xylostella* has developed into a major pest worldwide (Grzywacz et al., 2010). Following the misuse of insecticides in the field, *P. xylostella* has developed such severe resistance to most insecticides through multiple mechanisms (Endersby et al., 2011; Troczka et al., 2012). Although RNAi had been reported to be relatively inefficient in lepidoptera (Shukla et al., 2016; Zhu and Palli., 2020), RNAi for *P. xylostella* was proven to be feasible by injection or oral ingestion (Pan et al., 2017; Zhang et al., 2020). Thus, the control and resistance management of *P. xylostella* may be achieved by dsRNA ingestion and interference with specific target genes.

The ionotropic γ-aminobutyric acid (GABA) receptors (GABARs) are the major inhibitory receptors widely distributed in the central nervous system of insects (Ozoe., 2013). The RDL subunit encoded by the resistance to dieldrin (*Rdl*) gene is the target of non-competitive antagonists (NCAs) and allosteric modulators (Buckingham et al., 2017; Ozoe., 2021). *P. xylostella* possessed at least two *Rdl* genes (Yuan et al., 2010; Li et al., 2021). Our previous *Xenopus laevis* oocytes studies showed the inhibitory potencies of fipronil on *Px*RDL2 was about 40-fold lower than on *Px*RDL1. In addition, knockout of *PxRdl1* reduced the potencies of fipronil and pyrazoloquinazoline **5a**, while knockout of *PxRdl2* increased their potencies (Li et al., 2021; Li et al., 2023). These results indicate *PxRdl2* may play an important role in determining the susceptibility of *P. xylostella* to GABAR-targeting compounds, which is helpful to explore the toxicology and resistance to insecticides. Hence, we verified the effect of the knockout of *PxRdl* on the insecticidal activities of GABAR-targeting compounds and considered whether it is possible to develop *PxRdl*-based RNAi to enhance their insecticidal activities.

In this study, the *PxRdl2* dsRNA which expressed in HT115 *Escherichia coli* (named ds-B in this paper) successfully inhibited *PxRdl2* transcription, and significantly enhanced the insecticidal activities of the pyrazoloquinazolines and isoxazolines against both susceptible and fipronil-resistant *P. xylostella* strains. In addition, we also verified the effect of UV irradiation on ds-B to ensure that it could remain stable when exposed to sunlight in field applications. In summary, our study provides new insight for pest management, and may be beneficial for insecticide resistance management and reducing the usage of insecticides, which could be part of an integrated pest management (IPM) approach.

## 2. Materials and methods

### 2.1. Chemicals and insects

The pyrazoloquinazoline **5a** (IUPAC name: methyl 6-chloro-2-cyano-5-(2-methoxy-2-oxoethyl)-8-(trifluoromethyl)-3-((trifluoromethyl)sulfinyl)-4,5-dihydropyrazolo[1,5-α]quinazoline-5-carboxylate; CAS number: 2232118-39-5; purity > 98%. Jiang et al., 2020) and **4a** (UPAC name: diethyl 9-chloro-2-cyano-7-(trifluoromethyl)-3-((trifluoromethyl)sulfinyl)pyrazolo[1,5-α]quin azoline-5,5(4H)-dicarboxylate; purity > 98%. Yang et al., 2020) were synthesized in our laboratory. The standards of fipronil (IUPAC name: 5-amino-1-[2,6-dichloro-4-(trifluoromethyl)phenyl]-4-(trifluoromethylsulfinyl)pyrazo le-3-carbonitrile; CAS number: 120068-37-3; purity, 95%), fluxametamide (IUPAC name: 4-[5-(3,5-dichlorophenyl)-5-(trifluoromethyl)-4*H*-1,2-oxazol-3-yl]-*N*-[(*E*)-methoxyim inomethyl]-2-methylbenzamide; CAS number: 928783-29-3; purity, 95%), chlorantraniliprole (IUPAC name: 5-bromo-*N*-[4-chloro-2-methyl-6-(methylcarbamoyl)phenyl]-2-(3-chloropyridin-2-yl) pyrazole-3-carboxamide; CAS number: 500008-45-7; purity, 95%), fluralaner (IUPAC name: 4-[5-(3,5-dichlorophenyl)-5-(trifluoromethyl)-4*H*-1,2-oxazol-3-yl]-2-methyl-*N*-[2-oxo-2-(2,2,2-trifluoroethylamino)ethyl]benzamide; CAS number: 864731-61-3; purity, 95%) and abamectin (IUPAC name: (1’*R*,2*R*,3*S*,4’*S*,6*S*,8’*R*,10’*E*,12’*S*,13’*S*,14’*E*,16’*E*,20’*R*,21’*R*,24’*S*)-2-butan-2-yl-21’,24’-di hydroxy-12’-[(2*R*,4*S*,5*S*,6*S*)-5-[(2*S*,4*S*,5*S*,6*S*)-5-hydroxy-4-methoxy-6-methyloxan-2-yl]oxy-4-methoxy-6-methyloxan-2-yl]oxy-3,11’,13’,22’-tetramethylspiro[2,3-dihydrop yran-6,6’-3,7,19-trioxatetracyclo[15.6.1.1^4,8^.0^20,24^]pentacosa-10,14,16,22-tetraene]-2’-one;(1’*R*,2*R*,3*S*,4’*S*,6*S*,8’*R*,10’*E*,12’*S*,13’*S*,14’*E*,16’*E*,20’*R*,21’*R*,24’*S*)-21’,24’-dihydroxy-12’-[(2*R*,4*S*,5*S*,6*S*)-5-[(2*S*,4*S*,5*S*,6*S*)-5-hydroxy-4-methoxy-6-methyloxan-2-yl]oxy-4-methoxy-6-methyloxan-2-yl]oxy-3,11’,13’,22’-tetramethyl-2-propan-2-ylspiro[2,3-dih ydropyran-6,6’-3,7,19-trioxatetracyclo[15.6.1.1^4,8^.0^20,24^]pentacosa-10,14,16,22-tetraen e]-2’-one; CAS number: 71751-41-2; purity, 95%) were purchased from TargetMol (Shanghai, China). The other chemicals used in this research were purchased from Sigma-Aldrich except with special instructions.

The susceptible and fipronil-resistant (*Px*RDL1 carried A302S mutation) *P. xylostella* strains were generously provided by Dr. Minsheng You (Fujian Agriculture and Forestry University, China). The *Px*RDL homozygous knockout strains (*PxRdl1*KO and *PxRdl2*KO) were established previously (Li *et al*., 2021). All strains were maintained separately at 25±1°C and 65±5% relative humidity under 16:8 h (light: dark) photoperiod.

### 2.2. Bioassays

Leaf dip bioassays were used to evaluate the lethal concentrations (LCs) of compounds against the susceptible, *PxRdl*KO and fipronil-resistant *P. xylostella* strains (Li et al., 2021). The effect of *PxRdl* RNAi on the insecticidal activities of compounds were investigated by the feeding of leaves containing both dsRNA and compounds. The LC_50_ and LC_20_ concentrations of compounds were used for *PxRdl1* and *PxRdl2* RNAi, respectively. Mortality was recorded after 48 h of continuous ingestion of dsRNA and compounds. The theoretically reduced dose was calculated by the following formula: (Compound alone - Compound adding dsRNA) /Compound alone × 100%.

### 2.3. synthesis of PxRdl dsRNA

The full length of *PxRdl1* and *PxRdl2* had been cloned from the cDNA of the susceptible *P. xylostella* strain in the previous study (Li et al., 2021). A 428-bp fragment of *PxRdl1* and a 441-bp fragment of *PxRdl2* were amplified from the full length of *PxRdl1* and *PxRdl2*, respectively. The PCR products were used as templates to synthesize specific dsRNA using the *in vitro* Transcription T7 Kit (TaKaRa Biotechnology, Dalian, China) (dsRNA-*Vitro*, ds-V). The 289-bp fragment of *GFP* was amplified from the cloning vector pE*GFP*-N1-ro1*GFP* (Plasmid #82369, Addgene), and then, a similar method was applied to synthesize specific *GFP* dsRNA (named *GFP* ds-V).

The 289-bp *GFP*, 428-bp *PxRdl1* and 441-bp *PxRdl2* fragments were cloned into L4440 vector (Plasmid #1654, Addgene) and transformed into HT115 *E. coli* competent cells. The recombinant bacteria were induced by isopropyl *β*-D-1-thiogalactopyranoside to produce dsRNA. Subsequently, the recombinant bacteria were collected by centrifugation at 5000×g for 5 min at 4°C and suspended in nuclease-free water. Sonication treatment was carried out using a low-temperature, ultrahigh-pressure cell crusher (Guangzhou Juneng Nano & Biotechnology Co., Ltd, Guangzhou, China) at 1800 bar. The LB agar plates containing 100 μg mL^-1^ ampicillin were used to evaluate whether the bacteria were totally inactivated.

### 2.4. qRT-PCR absolute quantification of dsRNA

qRT-PCR was performed using SYBR^®^ Premix Ex Taq™ II (Takara Biotechnology) and CFX96 Connect Real-Time system (Bio-Rad Laboratories, Inc., Hercules, CA, USA). Based on the absolute quantification method (Rutledge and Cote, 2003), the quantity of dsRNA was confirmed by threshold (CT) values, which relate to an established standard curve. The standard curve for dsRNA was established by plotting the logarithm of a 10- to 10^5^-fold dilution of the starting solution of template cDNA with inserts against the corresponding CT values.

### 2.5. qRT-PCR relative quantification of PxRdl transcription

The ds-B (4 μg μL^-1^) or ds-V (4 μg μL^-1^) was applied to leaves by leaf dipping and then naturally air-dried. The early third-instar larvae were continuously fed on the leaves that contain dsRNA for 48 h. The gene-specific primers were designed to test a segment of the mRNA external to the segment targeted by the dsRNA. The *β-actin* (GenBank No. JN410820.1) was used as housekeeping gene and the data was analyzed using 2^−ΔΔCT^ method (Livak & Schmittgen, 2001). Each treatment used at least 10 living larvae. The controls received *GFP* ds-B (4 μg μL^-1^) and *GFP* ds-V (4 μg μL^-1^). All primers are listed in Tables S1.

### 2.6. UV irradiation of dsRNA

To better simulate the field condition, a UV irradiation experiment was used to assess the stability of dsRNA. The *PxRdl2* ds-B (4 μg μL^-1^), *PxRdl2* ds-V (4 μg μL^-1^), *GFP* ds-B (4 μg μL^-1^) and *GFP* ds-V (4 μg μL^-1^) were placed on the surface of glassware and continuously irradiated for 1 h, 2h and 4 h using a UV lamp (254 nm, 8 W), followed by fed to *P*. *xylostella* through leaves for gene expression analysis after 48 h.

### 2.7. Electrophysiological recording for X. laevis oocytes

The potencies of pyrazoloquinazoline **4a**, fluxametamide and fluralaner were recorded under two-electrode voltage clamp (TEVC) mode. The details of electrophysiological recordings were similar as previously described (Li et al., 2021; Li et al., 2023). Each experiment was performed on 5-6 different oocytes obtained from at least two frogs.

### 2.8. Statistical analysis

The LCs were calculated with SPSS 22.0 (SPSS Inc., Chicago, IL, USA) and analyzed using the regression-probit analysis. The bioassay data were analyzed by Student’s t test (\**p*< 0.05) using SPSS 22.0. The gene expression was analyzed by one-way ANOVA followed by Tukey test (*p*< 0.05) using SPSS 22.0. The 50% inhibitory concentrations (IC_50_s) and Hill coefficients (nHs) were obtained by nonlinear regression analysis using GraphPad Prism 8.0 (GraphPad Software Inc., San Diego, CA, USA).

## 3. Results

### 3.1. Insecticidal activities of compounds against PxRdl knockout P. xylostella

To test the roles of the two *PxRdl* homologous genes in the interaction with pyrazoloquinazolines and isoxazolines, the responses of the *PxRdl* knockout strains and susceptible strain were measured (Table 1 and Fig. 1). Knockout of *PxRdl1* significantly decreased **4a** susceptibility by 2.85-fold, whereas knockout of *PxRdl2* slightly but significantly increased **4a** susceptibility by 2.22-fold. Compare with the susceptible strain, the insecticidal activity of fluxametamide against the *PxRdl*1KO and *PxRdl*2KO strains was significantly increased by 1.88- and 1.85-fold, respectively. Unlike fluxametamide, knockout of *PxRdl1* had no significant effect on fluralaner susceptibility, but knockout of *PxRdl2* increased fluralaner susceptibility by 2.50-fold. No significant difference in insecticidal activities was observed against the three strains for abamectin and chlorantraniliprole.

**Fig. 1.**
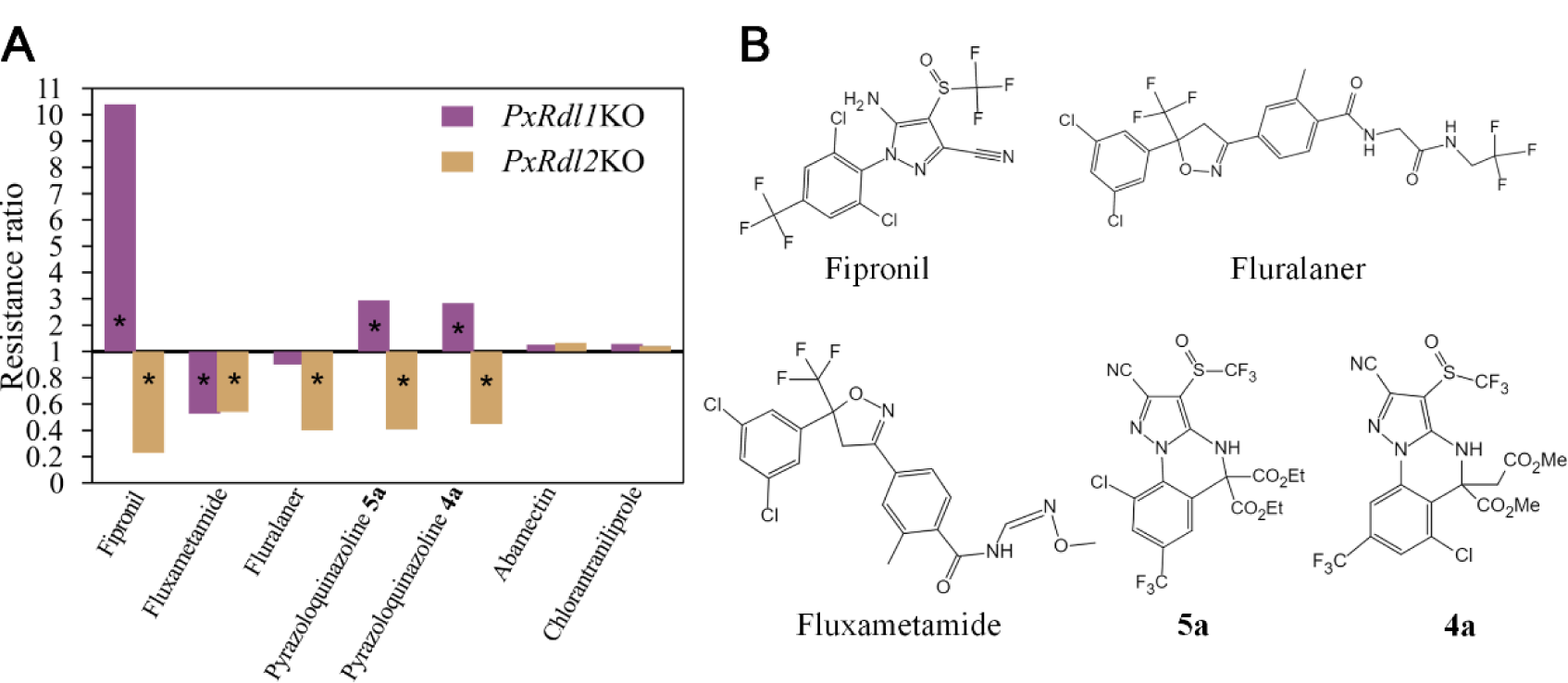
Differences in resistance levels. (A) Resistance levels of *PxRdl1*KO (*PxRdl1* knockout strain) and *PxRdl2*KO (*PxRdl2* knockout strain) to compounds compared with the susceptible strain of *Plutella xylostella*. Resistance ratio = LC_50_ value of *Rdl1*KO or *Rdl2*KO strain divided by LC_50_ value of the susceptible strain. The star indicates their LC_50_s are significantly different from those of susceptible individuals; (B) The structures of GABAR-targeting compounds.

**Table 1.** Insecticidal activities of compounds against third-instar *P. xylostella* larvae of susceptible, fipronil-resistant and *PxRdl* knockout strains.

| Compound | Strain | Slope ± SE | LC<sub>50</sub> (mg L<sup>-1</sup>)<br>(95% CI) | RR<sup>a</sup> |
| --- | --- | --- | --- | --- |
| Fipronil<sup>b</sup> | Susceptible | 2.66 ± 0.02 | 0.22 (0.19-0.26) | - |
| | *Rdl1*KO | 2.27 ± 0.22 | 2.29 (1.90-2.76) | 10.41 |
| | *Rdl2*KO | 2.58 ± 0.01 | 0.05 (0.04-0.06) | 0.23 |
| | Fipronil-resistant | 4.43 ± 0.48 | 41.38 (36.65-45.93) | 188.11 |
| **4a** | Susceptible | 3.01 ± 0.34 | 3.93 (3.41-4.60) | - |
| | *Rdl1*KO | 5.32 ± 0.58 | 11.19 (10.32-12.17) | 2.85 |
| | *Rdl2*KO | 2.50 ± 0.29 | 1.78 (1.51-2.11) | 0.45 |
| | Fipronil-resistant | 4.41 ± 0.50 | 5.80 (4.04-7.94) | 1.48 |
| **5a<sup>c</sup>** | Susceptible | 3.11 ± 0.37 | 2.23 (1.93-2.56) | - |
| | *Rdl1*KO | 2.43 ± 0.34 | 6.74 (6.03-7.53) | 3.02 |
| | *Rdl2*KO | 1.82 ± 0.14 | 0.84 (0.61-1.17) | 0.38 |
| | Fipronil-resistant | 3.19 ± 0.41 | 2.10 (1.83-2.38) | 0.94 |
| Fluxametamide | Susceptible | 2.65 ± 0.43 | 0.079 (0.051-0.59) | - |
| | *Rdl1*KO | 4.06 ± 0.44 | 0.042 (0.037-0.047) | 0.53 |
| | *Rdl2*KO | 3.21 ± 0.37 | 0.043 (0.038-0.050) | 0.54 |
| | Fipronil-resistant | 2.19 ± 0.21 | 0.061 (0.035-0.101) | 0.78 |
| Fluralaner | Susceptible | 4.40 ± 0.85 | 0.010 (0.009-0.012) | - |
| | *Rdl1*KO | 5.53 ± 0.73 | 0.009 (0.006-0.015) | 0.90 |
| | *Rdl2*KO | 4.67 ± 0.48 | 0.004 (0.003-0.005) | 0.40 |
| | Fipronil-resistant | 2.44 ± 0.31 | 0.013 (0.007-0.067) | 1.30 |
| Abamectin | Susceptible | 1.69 ± 0.24 | 0.012 (0.008-0.015) | - |
| | *Rdl1*KO | 2.30 ± 0.27 | 0.015 (0.012-0.018) | 1.25 |
| | *Rdl2*KO | 2.33 ± 0.25 | 0.016 (0.014-0.020) | 1.33 |
| Chlorantraniliprole | Susceptible | 1.30 ± 0.19 | 1.32 (0.88-1.81) | - |
| | *Rdl1*KO | 1.41 ± 0.19 | 1.70 (1.23-2.27) | 1.29 |
| | *Rdl2*KO | 1.11 ± 0.18 | 1.58 (1.03-2.25) | 1.20 |

<sup>a</sup>The resistance ratio was calculated by LC<sub>50</sub> values of *PxRdl1*KO, *PxRdl2*KO or resistant strains divided by LC<sub>50</sub> value of susceptible strain.
<sup>b</sup>The bioassay results of fipronil against the *P. xylostella* larvae of the susceptible, *PxRdl1*KO and *PxRdl2*KO strains were reported previously (Li et al., 2021).
<sup>c</sup>The bioassay results of **5a** against the *P. xylostella* larvae of the susceptible, *PxRdl1*KO and *PxRdl2*KO strains were reported previously (Li et al., 2023).

### 3.2. Production of bacteria expressing PxRdl dsRNA

The fragments of *GFP* and *PxRdls* were successfully transformed into HT115 *E. coli* cells (Fig. 2A), and the quantity of dsRNA was determined by qRT-PCR (Fig. 2B, 2C and 2D). For *GFP*, *PxRdl1* and *PxRdl2*, the PCR efficiencies which calculated according to the slope and the coefficient of correlation (*R*^2^) of the standard curve were 95.52%, 108.93% and 92.78%, respectively. The above results indicated the qRT-PCR was suitable for monitoring the content of dsRNA expressed in HT115 *E. coli* cells.

**Fig. 2.**
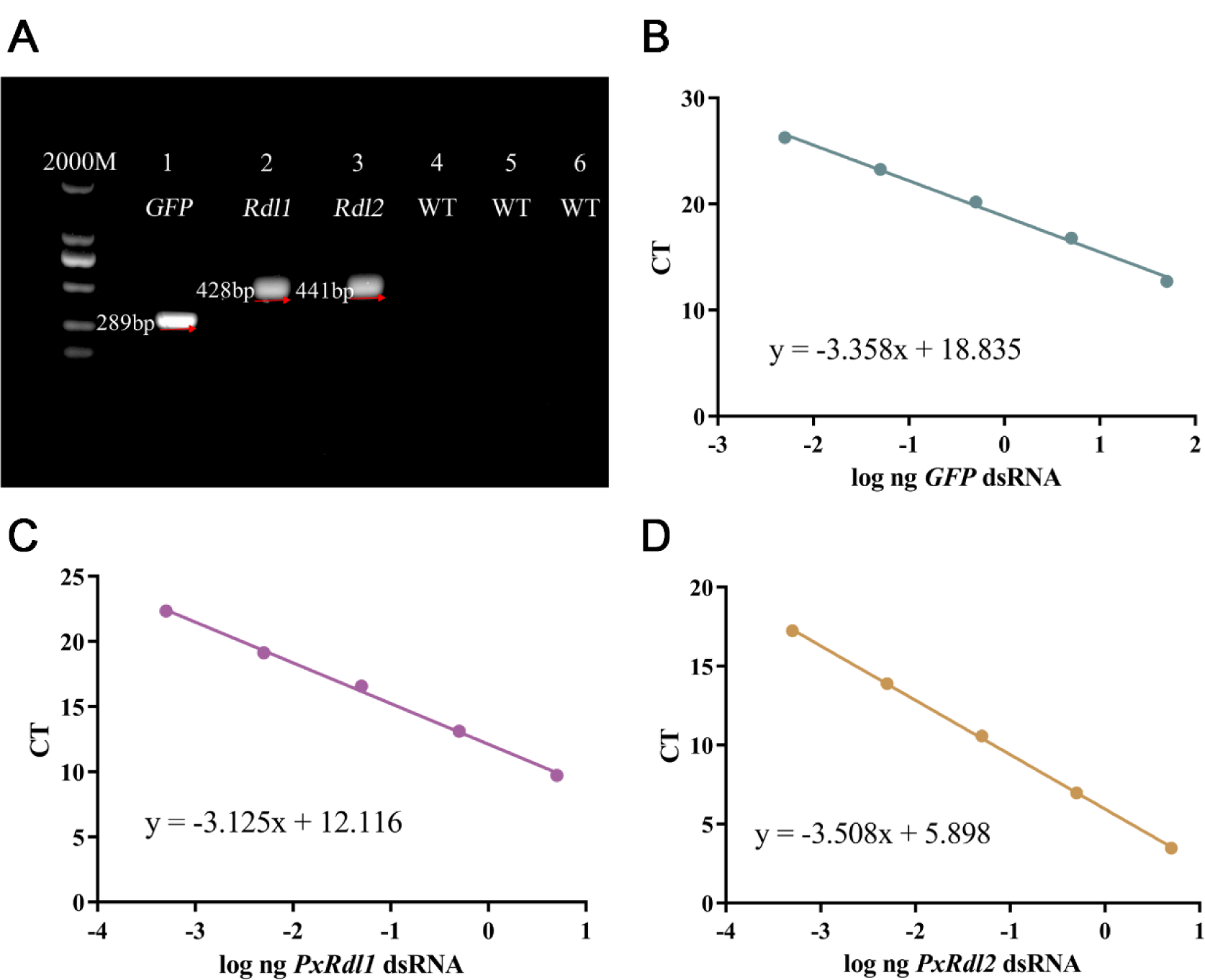
Production of HT115 *Escherichia coli* cells expressing dsRNA. (A) Expression of dsRNA by transformed HT115 *E. coli*; The specific primers used for the *GFP*, *PxRdl1* or *PxRdl2* genes produced amplicons of the expected fragments in transformed HT115 *E. coli* (lines 1, 2 and 3), whereas the same primers did not produce any fragment from non-transformed HT115 *E. coli* (WT) (lines 4, 5 and 6). (B), (C) and (D) Standard curves used for qRT-PCR absolute quantification of *GFP*, *PxRdl1* and *PxRdl2* dsRNA produced by *E. coli* suspensions.

### 3.3. PxRdl interference in P. xylostella

The *P. xylostella* larvae were continuously fed on the leaves coated with ds-B or ds-V for 48 h, and the transcript levels of *PxRdls* were determined by qRT-PCR. The results indicated the transcript levels of *PxRdls* were significantly inhibited by dsRNA treatment. Furthermore, ds-B provided a significantly higher interference efficiency compare with ds-V (*p* < 0.05) (Fig. 3A and 3B). The interference efficiency of each dsRNA was as follows: *PxRdl1* ds-B 63.63%; *PxRdl1* ds-V 35.37%; *PxRdl2* ds-B 64.61% and *PxRdl2* ds-V 38.44%.

**Fig. 3.**
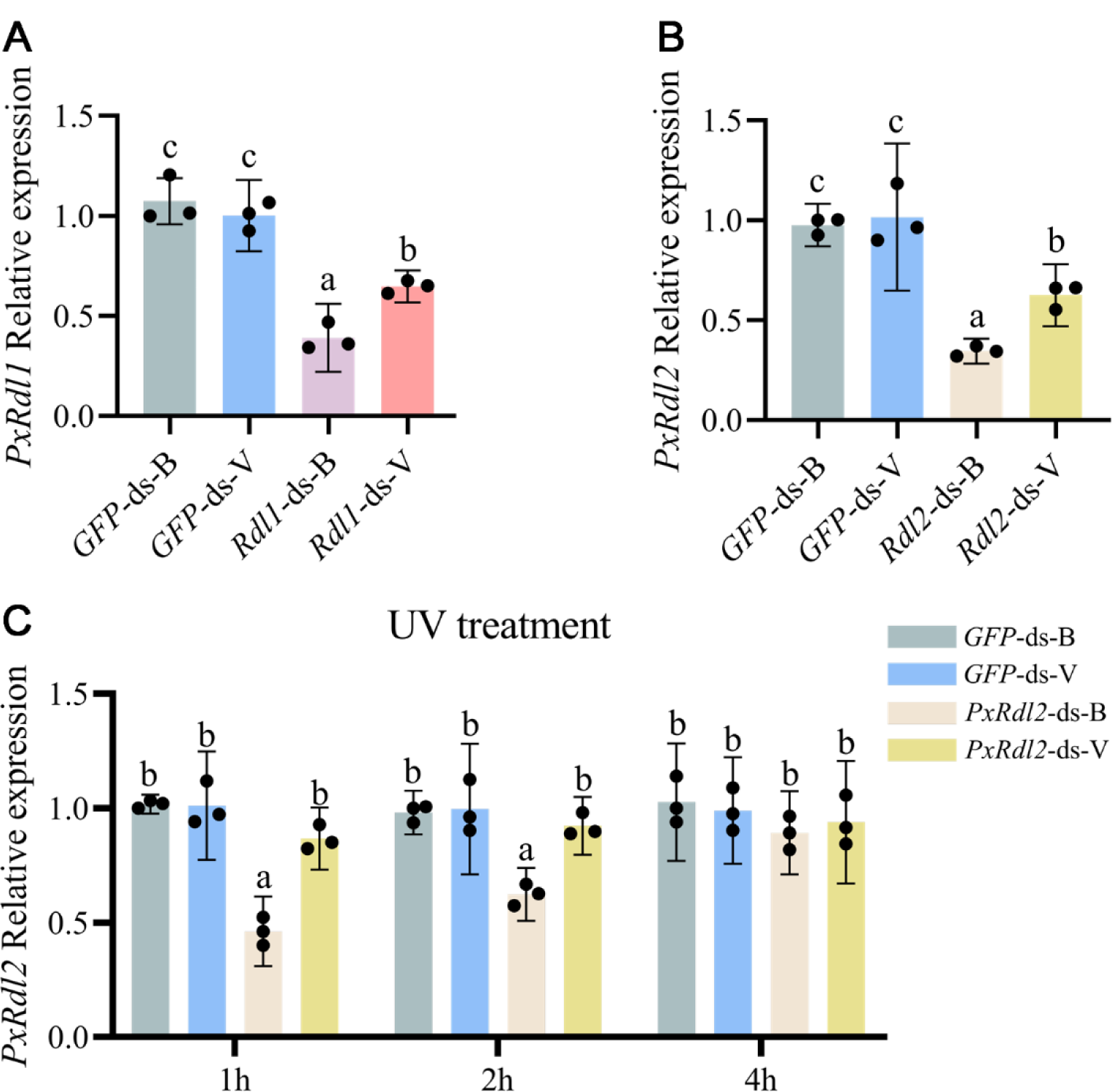
Effect of the dietary introduction of 4 μg μL^-1^ dsRNA on the relative *PxRdls* transcript levels of third-instar larvae of *Plutella xylostella*. (A) *PxRdl1*, (B) *PxRdl2*, (C) *PxRdl2* dsRNA treated with different UV irradiation durations (1 h, 2 h, and 4 h). ds-B: dsRNA expressed by HT115 *Escherichia coli*, ds-V: dsRNA synthesized *in vitro*. Lowercase letters (a b and c) indicate significance among different treatment groups (*p* < 0.05, Tukey test).

### 3.4. The effect of UV irradiation on PxRdl2 dsRNA

After *PxRdl2* ds-B or *PxRdl2* ds-V were continuously irradiated with UV for different durations, the treated dsRNA was fed to third-instar larvae through the leaves. The transcript level of *PxRdl2* was used to assess the stability of *PxRdl2* dsRNA (Fig. 3C). 2 h after UV irradiation, the *PxRdl2* ds-B still maintained high activity, and the interference efficiency of *PxRdl2* ds-B was significantly higher than that of other treatments. However, *PxRdl2* ds-V lost activity after 1 h of UV irradiation (*p* < 0.05). The interference efficiency of *PxRdl2* ds-B under different UV irradiation durations was 54.58% for 1 h and 36.53% for 2h, respectively (Fig. 3C).

### 3.5. Effect of PxRdl interference on larvae susceptibility to compounds

The LC_50_ concentrations of fipronil, pyrazoloquinazolines and isoxazolines were used for assessing the effect of *PxRdl1* interference on larvae susceptibility to compounds. After fipronil, **5a** and **4a** treatment, the mortalities of larvae of the susceptible *P. xylostella* strain in *PxRdl1* ds-B treatment groups significantly decreased by 20.56%, 25.00% and 28.60%, respectively, compare with that in GFP ds-B treatment groups. In contrast, the insecticidal activity of fluxametamide was significantly increased by adding *PxRdl1* ds-B, but the insecticidal activity of fluralaner didn’t change significantly (**Fig. 4a**).

**Fig. 4.**
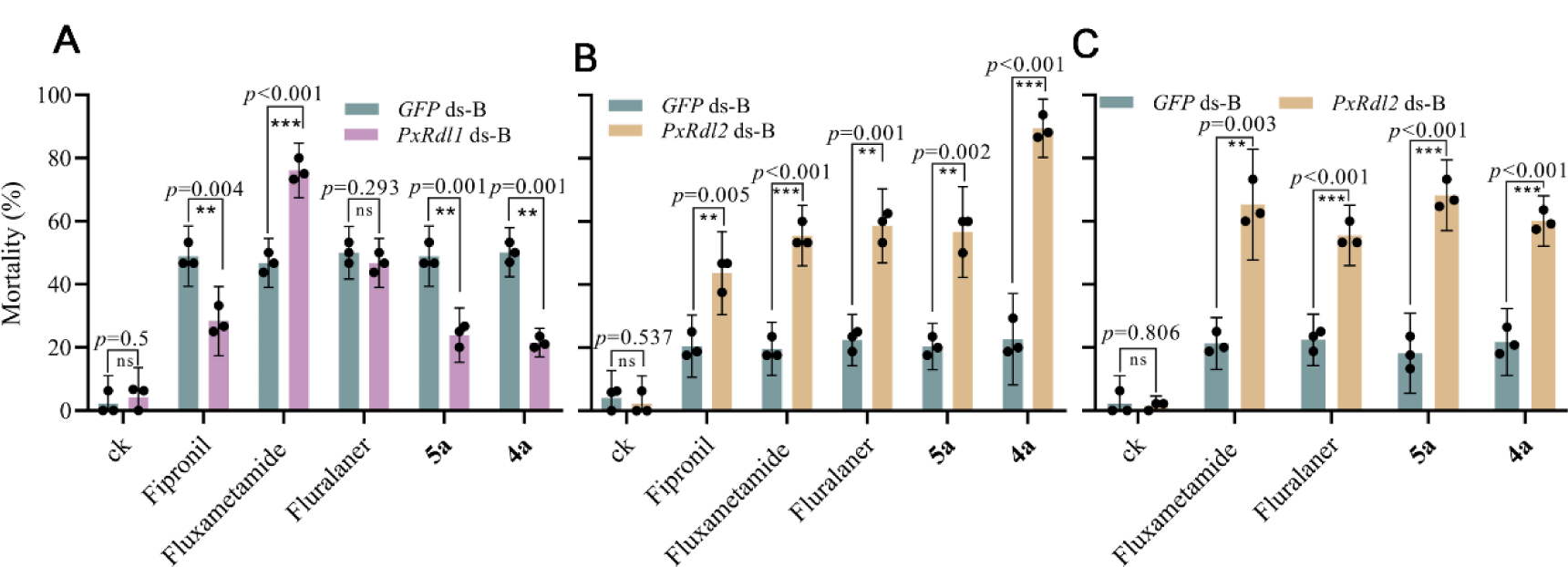
Effect of the dietary introduction of 4 μg μL^-1^ dsRNA and compound on the 48h mortality of third-instar larvae of *Plutella xylostella*. (A) *PxRdl1* dsRNA and compound against the susceptible strain, (B) *PxRdl2* dsRNA and compound against the susceptible strain, (C) *PxRdl2* dsRNA and compound against the fipronil-resistant strain. ds-B: dsRNA expressed by HT115 *Escherichia coli*, ds-V: dsRNA synthesized *in vitro*, CK: no treatment. The significantly different was calculated using Student’s t test (\**p* < 0.05, \*\**p* < 0.01, and \*\*\**p* < 0.001).

The bioassay results showed the suppression of *PxRdl2* transcription by RNAi followed by LC_20_ concentrations of compounds exposure increased the susceptibility of larvae of the susceptible *P. xylostella* strain to fipronil, **5a**, **4a**, fluxametamide and fluralaner, respectively, with 48 h mortalities of 43.61%, 56.67%, 89.55%, 55.55% and 58.61%, respectively (Fig. 4B). The effect of RNAi-mediated insecticidal activity enhancement was evaluated using the theoretical reduction in compound usage, and results showed *PxRdl2* RNAi reduced the doses required of five compounds by 38.89% to 80.25% (Table 2). Similarly, the insecticidal activities of **5a**, **4a**, fluxametamide, and fluralaner against fipronil-resistant *P. xylostella* were enhanced by adding of *PxRdl2* ds-B (Fig. 4C), with doses required of **5a**, **4a**, fluxametamide, and fluralaner reducing by 62.75%, 43.75%, 72.82% and 60.00%, respectively (Table 2).

**Table 2.** The theoretically reduced doses required of compounds against *Plutella xylostella* by adding *PxRdl2* dsRNA

| Compound | Strain | RNAi | Compound alone | Reduce dose |
| --- | --- | --- | --- | --- |
|  |  | LC<sub>20</sub> compound (mg L<sup>-1</sup>)<br>(95% CI) | (mg L<sup>-1</sup>) (95% CI) | (%)<sup>a</sup> |
| Fipronil | Susceptible | 0.11 (0.09-0.14) | 0.18 (0.15-0.21) | 38.89 |
| **4a** | Susceptible | 2.07 (1.66-2.44) | 10.48 (8.26-14.93) | 80.25 |
|  | Fipronil-resistant | 3.73 (1.57-4.99) | 6.61 (4.94-9.92) | 43.57 |
| **5a** | Susceptible | 0.94 (0.54-1.51) | 1.91 (1.04-3.83) | 50.79 |
|  | Fipronil-resistant | 2.85 (2.18-3.40) | 7.65 (6.69-9.14) | 62.75 |
| Fluxametamide | Susceptible | 0.038 (0.011-0.060) | 0.088 (0.056-0.190) | 56.82 |
|  | Fipronil-resistant | 0.025 (0.010-0.042) | 0.092 (0.056-0.166) | 72.82 |
| Fluralaner | Susceptible | 0.006 (0.005-0.007) | 0.011 (0.010-0.014) | 45.45 |
|  | Fipronil-resistant | 0.006 (0.005-0.010) | 0.015 (0.008-0.113) | 60.00 |

<sup>a</sup>Theoretically reduce dose = (Compound alone – compound adding dsRNA)/Compound alone × 100%.

### 3.6. Sensitivity of RDL GABAR to compounds

TEVC electrophysiology was performed in *Xenopus* oocytes to test the potencies of **4a**, fluxametamide and fluralaner on *Px*RDL GABAR (Fig. 5 and Table 3). **4a** showed inhibitory potency on *Px*RDL1 but not *Px*RDL2, with IC_50_ value of 2.02 *μ*mol/L (Fig. 5C). Fluralaner showed high inhibitory potency on *Px*RDL1 and *Px*RDL2, with IC_50_ values of 0.010 and 0.018 *μ*mol/L, respectively (Fig. 5B). Fluxametamide dose-response curves gave IC_50_ values of 0.020 *μ*mol/L for *Px*RDL1 and 0.017 *μ*mol/L for *Px*RDL2, respectively (**Fig. 5a**).

**Fig. 5.**
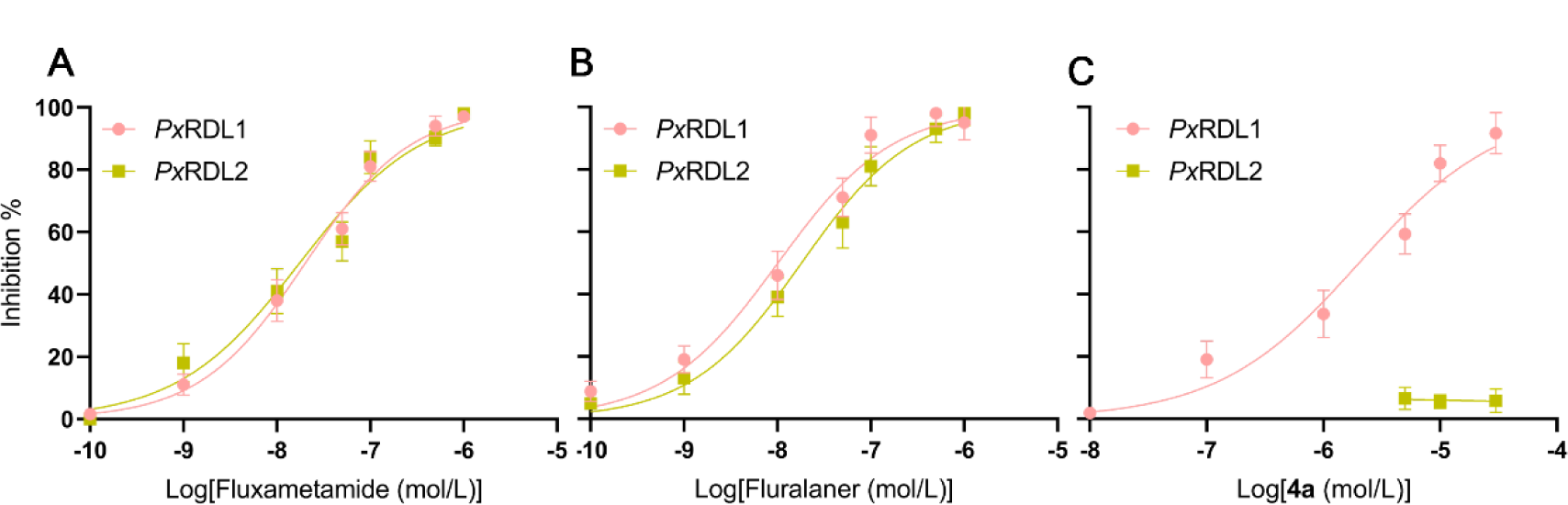
The inhibitory potencies of fluxametamide (A), fluralaner (B) and pyrazoloquinazoline **4a** (C) on *Px*RDLs GABAR. Each point represents the means ± SE of responses in 5-6 oocytes from at least two frogs.

**Table 3.** Potencies of antagonist in *Xenopus* oocytes expressed *Px*RDLs.

| Compounds | *PxRDL1* |  | *PxRDL2* |  |
| --- | --- | --- | --- | --- |
|  | IC<sub>50</sub> (95% CI) | Hill Slope | IC<sub>50</sub> (95% CI) | Hill Slope |
|  | (μmol/L) | (95% CI) | (μmol/L) | (95% CI) |
| **4a** | 2.02 (0.94-3.92) | 0.72 (0.44-1.27) | NI<sup>a</sup> | NI<sup>a</sup> |
| Fluxametamide | 0.020 (0.015-0.027) | 0.77 (0.62-0.96) | 0.017 (0.009-0.033) | 0.67 (0.46-1.05) |
| Fluralaner | 0.010 (0.006-0.017) | 0.71 (0.51-1.04) | 0.018 (0.012-0.025) | 0.73 (0.59-0.91) |

<sup>a</sup>NI= no inhibition at 30 μmol/L

## Discussion

In previous studies, a series of pyrazoloquinazoline derivatives were designed and synthesized via scaffold hopping based on the structure of fipronil. **5a** and **4a** both exhibited high insecticidal activities against *P. xylostella* (Jiang et al., 2020; Yang et al., 2020). **5a** could inhibit the function of *P. xylostella* GABAR, and the binding site of **5a** on *Px*RDL1 was similar to that of fipronil (Li et al., 2023). Our previous TEVC studies suggested the sensitivities of *Px*RDL2 to fipronil and **5a** were much lower than that of *Px*RDL1 (Li et al., 2021; Li et al., 2023). Similarly, **4a** also exhibited obvious inhibitory potency on *Px*RDL1 but not *Px*RDL2 (Fig. 5C). The different sensitivities of RDLs to fipronil or dieldrin are also found in *Varroa destructor* and *Chilo suppressalis* (Ménard et al., 2018; Sheng et al., 2018).

Knockout of *PxRdl1* significantly decreased the potencies of fipronil and two pyrazoloquinazolines, whereas knockout of *PxRdl2* significantly increased the potencies of three compounds (Table 1). This opposite trend was also observed in the cyclodienes against two *HaRdl* knockout *Helicoverpa armigera* strains (Wang et al., 2020). It is possible that the different pharmacological properties between *Px*RDL1 and *Px*RDL2 contribution to the sensitivities of *P. xylostella* to fipronil and two pyrazoloquinazolines, and *Px*RDL2 plays an antagonistic role in the action of the three compounds.

The fluxametamide and fluralaner are novel isooxazolines which allosteric inhibit insect GABAR by binding to a site different from the site for traditional NCAs (Gassel et al., 2014; Asahi et al., 2015; Asahi et al., 2018). There was no significant difference between the potencies of fluxametamide and fluralaner on *Px*RDL1 and *Px*RDL2 in TEVC studies. Consequently, knockout of *PxRdl2* both increased the insecticidal activities of fluxametamide and fluralaner. Taken together, our bioassay and TEVC results indicated *Px*RDL2 is involved in mediating the toxicities of GABAR-targeting compounds and may be an ideal site for dsRNA targeting.

Pest control based on RNAi, due to its specificity and minimal or nonexistent impact on nontarget species, provides a new opportunity for the development of sustainable IPM plans (Chen et al., 2021; Nandety et al., 2015). However, the reaction efficiency of lepidopteran insects to RNAi is uneven, and the dsRNA delivery mode will affect its interference efficiency. It has been reported that *H. armigera* and *Spodoptera littoralis* were sensitive to the orally delivered dsRNA (Caccia et al. 2020; Lim et al., 2016). Moreover, there were also reports of RNAi in *P. xylostella* using the orally delivered dsRNA (Mohamed et al., 2011; Guo et al., 2015; Chaitanya et al., 2017; Kang et al., 2022) which could provide us with references to control *P. xylostella* using RNAi.

The qRT-PCR results confirmed the transcript levels of *PxRdls* were inhibited by ds-B treatment (Fig. 3A and 3B). Although the knockout or knockdown of *PxRdl1* and *PxRdl2* alone didn’t affect the survival of *P. xylostella*, the bioassay results which performed on the *PxRdl* knockout strains allow for the extension of a new pest control method aimed at enhancing the insecticidal activities of GABAR-targeting compounds against *P. xylostella* through RNAi-mediated *PxRdl2* interference. However, the development of efficient, safety and economically sustainable RNAi delivery strategies is vital to achieve this goal.

In this study, the HT115 *E. coli* was used to product mass production of dsRNA targeting *PxRdls*. The qRT-PCR results confirmed the *PxRdl* dsRNA successfully interfered *PxRdl* transcription, and interference efficiency of ds-B was significantly higher than that of ds-V (Fig. 3A, Fig. 3B). Consist with the bioassay results which performed on the *PxRdl* knockout strains, the knockdown of *PxRdl2* which induced by ds-B also increased the insecticidal activities of fipronil, two pyrazoloquinazolines and two isooxazolines (Fig. 4B). In addition, the theoretically reduced doses of the compounds were calculated and ranged from 38.89% to 80.25% (Table 2). However, the *PxRdl1* interference only increased the mortalities of larvae in fluxametamide treatment (**Fig. 4a**). The above results further verified *PxRdl2* plays an important role in the response to the GABAR-targeting insecticides.

The fipronil-resistant *P. xylostella* strain which carried the A302S mutation in *Px*RDL1 subunit (He et al., 2012) showed about 188-fold resistance to fipronil (Table 1). However, the two isooxazolines and two pyrazoloquinazolines showed no cross resistance to fipronil (Table 1). Furthermore, the *PxRdl2* interference also increased the insecticidal activities of two isooxazolines and two pyrazoloquinazolines against the fipronil-resistant strain (Fig. 4C). The calculated theoretical reduction in the doses of the three compounds by 43.57% to 72.82% (Table 2). These results further confirmed *PxRdl2* maybe an ideal dsRNA target for using in fields to enhance the insecticidal activities of GABAR-targeting insecticides.

It was previously reported that dsRNA was protected by the bacterial shell and was not easily degraded in both the environment and the insect gut, which could allow for the dsRNA to exist/be released for a relatively long time (Lim et al., 2016; Vatanparast and Kim, 2017). Moreover, weakening the bacterial cell wall through sonication may stimulate dsRNA-uptake and enhance the efficiency of the induced RNAi response (Vatanparast and Kim, 2017; Vogel et al., 2019). In this study, the recombinant HT115 *E. coli* may also be more readily uptake by *P. xylostella* and cause higher inhibitory efficiency after sonication (Fig. 3A and 3B). The above studies provided support for the application of ds-B in the fields, where light is an important factor affecting the stability of dsRNA. As the intensity of UV irradiation in the laboratory is several times stronger than that of solar UV irradiation in the fields (San Miguel and Scott, 2016), a UV irradiation was carried out to assess the stability of ds-B. Compared with *in vitro* synthesized dsRNA, ds-B was stabler under UV irradiation (Fig. 3C).

In summary, oral *PxRdl2* ds-B combined with GABAR-targeting compounds could increase the insecticidal activities of the compounds against *P. xylostella*, and these combinations are suitable for both susceptible and fipronil-resistant strains. The stable ds-B can be prepared using bacteria in large quantities, which greatly reduces the associated costs. Therefore, a spray method which uses ds-B as a synergist of insecticides may be developed, which is beneficial for reducing insecticide doses, delaying pest resistance, and protecting nontarget organisms.

## Supporting information

Detailed information of primers used in this study.

## Acknowledgements

This work was supported by the Guangdong Modern Agricultural Industry Generic Key Technology Research and Development Innovation Team Project (Grant No. 2023KJ133).

## References

1. Al-Jamal, K.T., Nerl, H., Müller, K.H., Ali-Boucetta, H., Li, S.P., Haynes, P.D., Jinschek, J.R., Prato, M., Bianco, A., Kostarelos, K., Porter, A.E., 2011. Cellular uptake mechanisms of functionalised multi-walled carbon nanotubes by 3D electron tomography imaging. Nanoscale. 3, 2627–2635.

2. Asahi, M., Kobayashi, M., Matsui, H., Nakahira, K., 2015. Differential mechanisms of action of the novel *γ*-aminobutyric acid receptor antagonist ectoparasiticides fluralaner (A1443) and fipronil. Pest Manag. Sci. 71, 91–95.

3. Asahi, M., Kobayashi, M., Kagami, T., Nakahira, K., Furukawa, Y., Ozoe, Y., 2018. Fluxametamide: A novel isoxazoline insecticide that acts via distinctive antagonism of insect ligand-gated chloride channels. Pestic. Biochem. Physiol. 151, 67–72.

4. Bento, F.M., Marques, R.N., Campana, F.B., Demétrio, C.G., Leandro, R.A., Parra, J.R.P., Figueira, A., 2020. Gene silencing by RNAi via oral delivery of dsRNA by bacteria in the South American tomato pinworm, *Tuta absoluta*. Pest Manag. Sci. 76, 287–295.

5. Buckingham, S.D., Ihara, M., Sattelle, D.B., Matsuda, K., 2017. Mechanisms of action, resistance and toxicity of insecticides targeting GABA receptors. Curr. Med. Chem. 24, 2935–2945.

6. Caccia, S., Astarita, F., Barra, E., Di Lelio, I., Varricchio, P., Pennacchio, F., 2020. Enhancement of *Bacillus thuringiensis* toxicity by feeding *Spodoptera littoralis* larvae with bacteria expressing immune suppressive dsRNA. J. Pest Sci. 93, 303–314.

7. Chaitanya, B.N., Asokan, R., Sita, T., Rebijith, K.B., Ram Kumar, P., Krishna Kumar, N.K., 2017. Silencing of JHEH and EcR genes of *Plutella xylostella* (Lepidoptera: Plutellidae) through double stranded RNA oral delivery. J. Asia Pac. Entomol.,20, 637–643.

8. Chen, J.S., Peng, Y.C., Zhang, H.N., Wang, K.X., Zhao, C.Q., Zhu, G.H., Reddy Palli, S., Han, Z.J., 2021. Off-target effects of RNAi correlate with the mismatch rate between dsRNA and non-target mRNA. RNA Biol. 18, 1747–1759.

9. Chikami, Y., Kawaguchi, H., Suzuki, T., Yoshioka, H., Sato, Y., Yaginuma, T., Niimi, T., 2021. Oral RNAi of *diap1* results in rapid reduction of damage to potatoes in *Henosepilachna vigintioctopunctata*. J. Pest Sci. 94, 505–515.

10. Dhandapani, R.K., Gurusamy, D., Duan, J.J., Palli, S.R., 2020. RNAi for management of Asian long-horned beetle, Anoplophora glabripennis: identification of target genes. J. Pest Sci. 93, 823–832.

11. Endersby, N., Viduka, K., Baxter, S., Saw, J., Heckel, D., McKechnie, S., 2011. Widespread pyrethroid resistance in Australian diamondback moth, *Plutella xylostella* (L.), is related to multiple mutations in the para sodium channel gene. Bull. Entomol. Res. 101, 393–405.

12. Fire, A., Xu, S.Q., Montgomery, M.K., Kostas, S.A., Driver, S.E., Mello, C.C., 1998. Potent and specific genetic interference by double-stranded RNA in *Caenorhabditis elegans*. Nature. 391, 806–811.

13. Gassel, M., Wolf, C., Noack, S., Williams, H., Ilg, T., 2014. The novel isoxazoline ectoparasiticide fluralaner: selective inhibition of arthropod *γ*-aminobutyric acid-and L-glutamate-gated chloride channels and insecticidal/acaricidal activity. Insect Biochem. Mol. Biol. 45, 111–124.

14. Grzywacz, D., Rossbach, A., Rauf, A., Russell, D.A., Srinivasan, R., Shelton, A.M., 2010. Current control methods for diamondback moth and other brassica insect pests and the prospects for improved management with lepidopteran-resistant Bt vegetable brassicas in Asia and Africa. Crop. Prot. 29, 68–79.

15. Guo, Z.J., Kang, S., Zhu, X., Xia, J.X., Wu, Q.J. and Wang, S.L. Xie, W., Zhang, Y.J., 2015. The novel ABC transporter ABCH1 is a potential target for RNAi-based insect pest control and resistance management. Sci. Rep. 5, 1–14.

16. He, W.Y, You, M.S, Vasseur, L., Yang, G., Xie, M., Cui, K., Bai, J.L., Liu, C.H., Li, X.J., Xu, X.F., H, S.G., 2012. Developmental and insecticide-resistant insights from the de novo assembled transcriptome of the diamondback moth, *Plutella xylostella*. Genomics, 99, 169–177.

17. Hernández-Soto, A., Chacón-Cerdas, R., 2021. RNAi crop protection advances. Int. J. Mol. Sci. 22, 12148.

18. Jiang, X.Y., Yang, S., Yan, Y., Lin, F., Zhang, L., Zhao, W.J., Zhao, C., Xu, H.H., 2020. Design, Synthesis, and Insecticidal Activity of 5,5-Disubstituted 4, 5-Dihydropyrazolo [1,5-*α*] quinazolines as Novel Antagonists of GABA Receptors. J. Agric. Food Chem. 68, 15005–15014.

19. Kang, S., Sun, D., Qin, J.Y., Guo, L., Zhu, L.H., Bai, Y., Wu, Q.J., Wang, S.L., Zhou, X.G., Guo, Z.J., Zhang, Y.J., 2022. Fused: a promising molecular target for an RNAi-based strategy to manage Bt resistance in *Plutella xylostella* (L.). J. Pest Sci. 95, 101–114.

20. Koch, A., Kogel, K.H., 2014. New wind in the sails: improving the agronomic value of crop plants through RNAi-mediated gene silencing. Plant Biotechnol. J. 12, 821–831.

21. Li, B.J., Wang, K.K., Chen, D.P., Yan, Y., Cai, X.L., Chen, H.M., Dong, K., Lin, F., Xu, H.H., 2021. Distinct roles of two RDL GABA receptors in fipronil action in the diamondback moth (*Plutella xylostella*). Insect Sci. 28, 1721–1733.

22. Li, B.J., Yan, Y., Yao, G.K., Zhang, L., Lin, F., Xu, H.H., 2023. Mode of Action of Novel Pyrazoloquinazoline on Diamondback Moth (*Plutella xylostella*) Ligand-gated Chloride Channels. J. Agric. Food Chem. doi: 10.1021/acs.jafc.3c01270.

23. Lim, Z. X., Robinson, K. E., Jain, R. G., Sharath Chandra, G., Asokan, R., Asgari, S., Mitter, N., 2016. Diet-delivered RNAi in *Helicoverpa armigera* − Progresses and challenges. J. Insect Physiol. 85, 86–93.

24. Lu, Y., Deng, X., Zhu, Q., Wu, D., Zhong, J., Wen, L., Yu, X., 2023. The dsRNA Delivery, Targeting and Application in Pest Control. Agronomy. 13, 714.

25. Ménard, C., Folacci, M., Brunello, L., Charreton, M., Collet, C., Mary, R., Rousset, M., Thibaud, J.-B., Vignes, M., Charnet, P., Cens, T., 2018. Multiple combinations of RDL subunits diversify the repertoire of GABA receptors in the honey bee parasite *Varroa destructor*. J. Biol. Chem. 293, 19012–19024.

26. Mohamed, A.A.M., Kim, Y., 2011. A target-specific feeding toxicity of *β1* integrin dsRNA against diamondback moth, *Plutella xylostella*. Arch. Insect Biochem. Physiol. 78, 216–230.

27. Nandety, R.S., Kuo, Y.W., Nouri, S., Falk, B.W., 2015. Emerging strategies for RNA interference (RNAi) applications in insects. Bioengineered. 6, 8–19.

28. Ozoe, Y., 2013. Chapter four – *γ*-Aminobutyrate- and glutamate-gated chloride channels as targets of insecticides. In: Cohen, E. (Ed.), Advances in Insect Physiology. Academic Press, The Boulevard, Langford Lane, Kidlington, Oxford, OX51GB, UK, pp. 211–286.

29. Ozoe, Y., 2021. Ion channels and G protein-coupled receptors as targets for invertebrate pest control: From past challenges to practical insecticides. Biosci. Biotechnol. Biochem. 85, 1563–1571.

30. Pan, Z.Z., Xu, L., Liu, B., Zhang, J., Chen, Z., Chen, Q.X., Zhu, Y.J., 2017. PxAPN5 serves as a functional receptor of Cry2Ab in *Plutella xylostella* (L.) and its binding domain analysis. Insect Biochem. Mol. Biol. 105, 516–521.

31. Rutledge, R.G, Côté, C., 2003. Mathematics of quantitative kinetic PCR and the application of standard curves. Nucleic Acids Res. 31, 93e.

32. San Miguel, K., Scott, J.G., 2016. The next generation of insecticides: dsRNA is stable as a foliar-applied insecticide. Pest Manag. Sci. 72, 801–809.

33. Sheng, C.W., Jia, Z.Q., Ozoe, Y., Huang, Q.T., Han, Z.J., Zhao, C.Q., 2018. Molecular cloning, spatiotemporal and functional expression of GABA receptor subunits RDL1 and RDL2 of the rice stem borer *Chilo suppressalis*. Insect Biochem. Mol. Biol. 94, 18–27.

34. Shukla, J.N., Kalsi, M., Sethi, A., Narva, K.E., Fishilevich, E., Singh, S., Mogilicherla, K., Subba Reddy Palli, S.R., 2016. Reduced stability and intracellular transport of dsRNA contribute to poor RNAi response in lepidopteran insects. RNA Biol. 13, 656–669.

35. Sun, Y.J., Wang, P.P., Abouzaid, M., Zhou, H., Liu, H. and Yang, P., Lin, Y.J., Hull, J.J., Ma, W.H., 2020. Nanomaterial-wrapped dsCYP15C1, a potential RNAi-based strategy for pest control against *Chilo suppressalis*. Pest Manag. Sci. 76, 2483–2489.

36. Timmons, L., Court, D.L., Fire, A., 2001. Ingestion of bacterially expressed dsRNAs can produce specific and potent genetic interference in *Caenorhabditis elegans*. Gene. 263, 103–112.

37. Troczka, B., Zimmer, C.T., Elias, J., Schorn, C., Bass, C., Emyr Davies, T.G., Field, L.M., Williamson, M.S., Slater, R., Nauen, R., 2012. Resistance to diamide insecticides in diamondback moth, *Plutella xylostella* (Lepidoptera: Plutellidae) is associated with a mutation in the membrane-spanning domain of the ryanodine receptor. Insect Biochem. Mol. Biol. 42, 873–880.

38. Vatanparast, M., Kim, Y., 2017. Optimization of recombinant bacteria expressing dsRNA to enhance insecticidal activity against a lepidopteran insect, *Spodoptera exigua*. PloS One. 12, e0183054.

39. Vogel, E., Santos, D., Mingels, L., Verdonckt, T.W., Broeck, J.V., 2019. RNA interference in insects: protecting beneficials and controlling pests. Front. Physiol. 9, 1912.

40. Wang, J., Zhao, X.F., Yan, R., Wu, S.W., Wu, Y.D., Yang, Y.H., 2020. Reverse genetics reveals contrary effects of two *Rdl*-homologous GABA receptors of *Helicoverpa armigera* on the toxicity of cyclodiene insecticides. Pestic. Biochem. Physiol. 170, 104699.

41. Yang, S., Lai, Q.Q., Lai, F.W., Jiang, X.Y., Zhao, C., Xu, H.H., 2020. Design, synthesis, and insecticidal activities of novel 5-substituted 4,5-dihydropyrazolo [1,5-*α*] quinazoline derivatives. Pest Manag. Sci. 77, 1013–1022.

42. Yuan, G.R., Gao, W.Y., Yang, Y.H., Wu, Y.D., 2010. Molecular cloning, genomic structure, and genetic mapping of two *Rdl*-orthologous genes of GABA receptors in the diamondback moth, *Plutella xylostella*. Arch. Insect Biochem. Physiol. 74, 81–90.

43. Zhang, Z.T., Kong, J.R., De Mandal, S., Li, S.Z., Zheng, Z.H., Jin, F.L., Xu, X.X., 2020. An immune-responsive PGRP-S1 regulates the expression of antibacterial peptide genes in diamondback moth, *Plutella xylostella* (L.). Int. J. Biol. Macromol. 142, 114–124.

44. Zhu, K.Y., Palli, S.R., 2020. Mechanisms, applications, and challenges of insect RNA interference. Annu. Rev. Entomol. 65, 293–311.

45. Zotti, M., Dos Santos, E.A., Cagliari, D., Christiaens, O., Taning, C.N.T., Smagghe, G., 2018. RNA interference technology in crop protection against arthropod pests, pathogens and nematodes. Pest Manag. Sci. 74, 1239–1250.

