## Supplementary material for "Bacterial *PxRdl2* dsRNA increased the insecticidal activities of GABAR-targeting compounds against *Plutella xylostella*": Detailed information of primers used in this study.

**^*^Corresponding authors：**

| Primer name | Nucleotide sequence (5’-3’) | Function |
| --- | --- | --- |
| *GFP*-F | CAGTGCTTCAGCCGCTACCC | Amplification of *GFP* |
| *GFP*-R | AGTTCACCTTGATGCCGTTCTT |  |
| *PxRdl1*-F | AGCAATACTGGATTCATT | Amplification of *PxRdl1* |
| *PxRdl1*-R | TCTCTATATGGCACAGTTGT |  |
| *PxRdl2*-F | TGGTCTTCGCTAGTCTTT | Amplification of *PxRdl2* |
| *PxRdl2*-R | AATTTGTTGATTCCCTTC |  |
| T7-*GFP*-F | TAATACGACTCACTATAGGGCAGTGCTTCAGCCGCTACCC | *In vitro* synthesis of *GFP* dsRNA |
| T7-*GFP*-R | TAATACGACTCACTATAGGGAGTTCACCTTGATGCCGTTCTT |  |
| T7-*PxRdl1*-F | TAATACGACTCACTATAGGGAGCAATACTGGATTC | *In vitro* synthesis of *PxRdl1* dsRNA |
| T7-*PxRdl1*-R | TAATACGACTCACTATAGGGTCTCTATATGGCACA |  |
| T7-*PxRdl2*-F | TAATACGACTCACTATAGGGTGGTCTTCGCTAGTCT | *In vitro* synthesis of *PxRdl2* dsRNA |
| T7-*PxRdl2*-R | TAATACGACTCACTATAGGGATTTGTTGATTCCCTTCG |  |
| Re-*GFP*-F | AATTGGGTACCGGGCCCCCCCAGTGCTTCAGCCGCTACCC | Homologous recombination |
| Re-*GFP*-R | CTGATATCATCGATGAATTCAGTTCACCTTGATGCCGTTCTT |  |
| Re-*PxRdl1*-F | AATTGGGTACCGGGCCCCCCAGCAATACTGGATTCATT |  |
| Re-*PxRdl1*-R | CTGATATCATCGATGAATTCTCTCTATATGGCACAGTTGT |  |
| Re-*PxRdl2*-F | AATTGGGTACCGGGCCCCCCTGGTCTTCGCTAGTCT |  |
| Re-*PxRdl2*-R | CTGATATCATCGATGAATTCATTTGTTGATTCCCTTCG |  |
| qPCR-AQ-*GFP*-F | GCCGCTACCCCGACCACAT | Absolute quantification of dsRNA produced by the transformed bacteria |
| qPCR-AQ-*GFP*-R | CGCCCTCGAACTTCACCTC |  |
| qPCR-AQ-*PxRdl1*-F | GCGTAGAGACATTATCAGTT |  |
| qPCR-AQ-*PxRdl1*-R | AATTCGTTGCTCGTAGTAG |  |
| qPCR-AQ-*PxRdl2*-F | GGCTGCTGAGAAGAAA |  |
| qPCR-AQ-*PxRdl2*-R | GGGAGTGTGGGACCTA |  |
| qPCR-*Actin*-F | TGGCACCACACCTTCTAC | Relative quantification of *PxRdls* transcription |
| qPCR-*Actin*-R | CATGATCTGGGTCATCTTCT |  |
| qPCR-*PxRdl1*-F | GAAGTTGCCTCCAGACTGC |  |
| qPCR-*PxRdl1*-R | CCACCCTTTGAATGTGCC |  |
| qPCR-*PxRdl2*-F | GTCAGTCAGCTACGACAAACG |  |
| qPCR-*PxRdl2*-R | AAATCCAGGGTAAAATCCAT |  |

**Table S1.** Detailed information of primers used in this study.
